# Control of Thiol-Maleimide Reaction Kinetics in PEG Hydrogel Networks

**DOI:** 10.1101/198135

**Authors:** Lauren E. Jansen, Lenny J. Negrón-Piñeiro, Sualyneth Galarza, Shelly R. Peyton

## Abstract

Michael-type addition reactions are widely used to polymerize biocompatible hydrogels. The thiol-maleimide modality achieves the highest macromer coupling efficiency of the reported Michael-type pairs, but the resulting hydrogel networks are heterogeneous, because polymerization is faster than the individual components can be manually mixed. The reactivity of the thiol dictates the overall reaction speed, which can be slowed in organic solvents and acidic buffers. Since these modifications also reduce the biocompatibility of resulting hydrogels, we investigated a series of biocompatible buffers and crosslinkers to decelerate gelation while maintaining high cell viability. We found that lowering the polymer weight percentage (wt%), buffer concentration, and pH slowed gelation kinetics, but crosslinking with an electronegative peptide was optimal for both kinetics and cell viability. Slowing the speed of polymerization resulted in more uniform hydrogels, both in terms of visual inspection and the diffusion of small molecules through the network. However, reactions that were too slow resulted in non-uniform particle dispersion due to settling, thus there is a trade-off in hydrogel network uniformity versus cell distribution in the hydrogels when using these networks in cell applications.

## 1. Introduction

The high water content and bulk elasticity provided by hydrogels mimic the physical properties of many tissues [1]. For this reason, hydrogels offer new opportunities in regenerative medicine and tissue engineering to study and regulate cell behavior and function [2]. For example, the biophysical and biochemical properties of hydrogels can guide cell migration and promote proliferation *in vitro* [3,4]. The ability to guide cell behavior also allows hydrogels to be used therapeutically to restore function and structure to damaged organs [5,6]. Here we focus on a class of synthetic, biocompatible hydrogels, which can be modified with cell-attachment and -degradable motifs to provide bio-functionality [2,7].

Polyethylene glycol (PEG) is one of the most commonly used macromers for tissue engineering, because it is hydrophilic, has low non-specific adsorption of proteins, and is not degradable by mammalian enzymes [8]. PEG is easy to chemically functionalize because the basic structure is capped with hydroxyl end groups, enabling a variety of chemical reactions, such as photopolymerization, Micheal addition, and click, for hydrogel polymerization [9]. Of particular interest to us is the thiol-ene chemistry, which is commonly used to form PEG hydrogels because thiols react with numerous alkene groups under many conditions [8,10,11]. Additionally, biologically active peptides are easily incorporated into hydrogel matrices via the thiol in the amino acid cysteine. The Michael-type addition reaction is advantageous for cell encapsulation because thiols and alkenes crosslink under physiological conditions, with no by-products, and no need for free radical initiating chemicals [12,13].

The maleimide functional group is the most efficient base-catalyzed Michael-type acceptor, avoiding toxic chemicals and/or light-mediated crosslinking that can both limit cell viability [8]. The high reaction efficiency results in gels with broader stiffness ranges compared to other functional groups, but it also causes the polymer network to assemble faster than individual components can be uniformly mixed [8,14], resulting in heterogeneous networks. In tissue engineering, this gives the advantage of forming hydrogels *in situ* almost immediately [15]; however, this leads to non-uniform ligand densities and crosslinking gradients [14,16], which can confound results when linking cell behaviors to stiffness cues. Although mostly overlooked by the field, two key studies have begun to address the kinetics of these Michael-type addition reactions, with the goal of finding new ways to control or slow the crosslinking speed of these networks to achieve more uniform gels [13,14]. Their main suggestion is to change the pKa of the thiol, or change the buffer/solvent conditions to interfere with thiolate formation, all of which slow the reaction [14,17-19]. Lowering pH and changing buffer conditions are easy to implement and can slow the gelation kinetics, but the pH range tolerated by cells is rather limited.

We attempt to resolve this by reporting a broad range of buffers that are both appropriate for cell culture and reduce the speed of the thiol-maleimide reaction in a PEG system. We measured the polymerization rate as a function of macromer concentration, buffer pH and strength, the presence of an external catalytic base, and the pKa of the Michael-type donor. The kinetics of the thiol-maleimide reaction are significantly slowed, to different degrees, using each of these approaches. Increasing the thiol pKa by incorporating a negatively charged amino acid (particularly glutamate) in the peptide crosslinker was optimal for maintaining cell viability and decreasing the reaction speed. However, these crosslinkers do need to be synthesized, which adds an additional level of cost, time, and complexity to the hydrogel synthesis process. Using 0.1x PBS at low pH, while supplementing the reaction buffer with high glucose, also slowed the reaction while maintaining cell viability. Though this reduction in gelation kinetics was not as significant as the effect with electronegative crosslinkers, it is significantly cheaper and more applicable to a broad range of crosslinkers. While slowing the reaction improved user handling, we also found that too slow of a reaction, particularly with the glutamate functionalized crosslinkers, led to non-uniform bead distribution, highlighting the tradeoff between the different approaches. Overall, slowing the crosslinking reaction speed, while maintaining high cell viability, improves the users ability to form PEG hydrogels and increases its applications in tissue engineering and as bench-top hypothesis test-beds.

## 2. Materials and Methods

### 2.1 Buffer preparation

Buffer solutions were prepared in nanopure water as follows. 10x Phosphate buffered saline (PBS) contained 1370 mM NaCl, 27 mM KCl, 80 mM Na_2_HPO_4_, and 20 mM KH_2_PO_4_ (Thermo Fisher Scientific, Waltham, MA). 10x Citrate buffer contained 100 mM sodium citrate dihydrate (Sigma-Aldrich, St. Louis, MO), 1370 mM NaCl, and 27 mM KCl (Thermo). Dulbecco’s Modified Eagle Medium (DMEM) supplement was added at 13.37 g/L, according to manufacturer’s instructions. 2 mM triethanolamine (TEOA) was prepared in 1x PBS. The pH was measured using an Orion Star A111 pH meter (Thermo) and adjusted with 1 M NaOH or HCl after the addition of all chemicals and/or supplements.

### 2.2 Formation of 3D hydrogels

Hydrogels were prepared with a 4-arm PEG-maleimide (P4M, Average Mn 20 kDa, >95% purity, JenKem, Plano, TX) crosslinked with one of the following dithiol terminated macromers: PEG-dithiol (PDT, Mn 1000, >95% purity, Sigma-Aldrich), CRG (GCRGIPESLRAGGRC), CRE (GCREIPESLRAGERC), or CRD with a number of different peptide sequences (GCRDIPESLRAGDRCG, GCRDPQGIWGQDRCG, GCRDVPLSLYSGDRCG, GCRDGPLGLWARDRCG)[20,21]. P4M was dissolved in the stated buffer and mixed with PDT at a molar ratio of 1:1 thiol to maleimide in a 10µL total volume. Gelation time was recorded as the initial point of polymer mixing to the time where no further mixing could be done by hand with a pipet. All gelation points were normalized to an internal control 10 wt% hydrogel in 1x PBS pH 7.0, which polymerized in roughly 3 seconds.

### 2.3 Peptide synthesis

Peptides were synthesized on a CEM’s Liberty Blue automated solid phase peptide synthesizer (CEM, Matthews, NC) using Fmoc protected amino acids (Iris Biotech GMBH, Germany). Peptide was cleaved from the resin by sparging nitrogen gas in trifluoroacetic acid (triisopropylsilane: water: 2,2′-(Ethylenedioxy)diethanethiol 92.5:2.5:2.5:2.5 % by volume, Sigma-Aldrich) for 3 hours at 25°C in a 100 mL reaction vessel (ChemGlass). Resin was filtered, and the peptide was precipitated using -80°C diethyl-ether (Thermo). Molecular weight was validated using a MicroFlex MALDI-TOF (Bruker, Billerica, MA) in a α-cyano-4-hydroxycinnamic acid matrix. Peptides were purified to ≥95% on a VYDAC reversed-phase c18 column attached to a Waters 2487 dual λ Absorbance Detector and 1525 binary HPLC pump (Waters, Milford, MA). The following peptides were synthesized: GCRGIPESLRAGGRC, GCREIPESLRAGERC, GCRDVPLSLYSGDRCG, GCRDGPLGLWARDRCG, and the following sequences were purchased from GenScript (Piscataway, NJ) at >95% purity: GCRDIPESLRAGDRCG, GCRDPQGIWGQDRCG.

### 2.4 Fiberoptic pH microsensor

Monitoring pH of PEG scaffolds was adapted from a previous protocol[22]. The needle-type pH microsensors (PreSens, Germany) were calibrated to the pH of the polymer buffer solvent in accordance to the manufacturer’s instructions. The probe was placed in the appropriate buffer for 30 minutes before measurements were made. The sensor was fully immersed into either precursor hydrogel solutions or a 20µL hydrogel, that had been polymerized 30 minutes prior, at 25°C. Measurements made at intervals of 5 seconds for a duration of 1 minute were averaged for a final pH value.

### 2.5 Hydrogel mechanical characterization

Indentation testing was performed on 10µL hydrogels after gelation and swelling in PBS for 48 hours in a custom-built instrument previously described[1,23]. Here, we brought a 0.75 mm cylindrical flat steel probe in contact with the top of the sample, which kept the sample height (h) to contact radius (a) between 0.5<a/h<2. A maximum force of 2mN was applied to hydrogels at a fixed displacement rate (20µm/s). Material compliance was analyzed using a correction ratio between the contact radius and the sample height to account for the dimensional confinement, as previously described [24].

### 2.6 LIVE/DEAD stain for cell viability

All cell lines were cultured at 37 °C and 5% CO 2. Cell culture supplies were purchased from Thermo with the exception of bovine insulin (Sigma-Aldrich). The cell lines LNCaPcol, PC-3, and SKOV-3, were cultured in Roswell Park Memorial Institute (RPMI) medium supplemented with 10% fetal bovine serum (FBS) and 1% penicillin/ streptomycin (Pen/Strep). OVCAR-3 cells were cultured in RPMI with 20% FBS, 1% Pen/Strep, and 0.01 mg/mL bovine insulin. MDA-MB-231, BT549, and SkBr3 breast cancer cells were cultured in DMEM with 10% FBS and 1% Pen/Strep. Cells were encapsulated at 3.0x10^5^ cells/mL in 1x PBS or 1x citrate with and without DMEM supplemented in the buffer solution, see methods above. Cell viability was determined using a LIVE/DEAD stain 2 hours post-encapsulation. Fluorescent images were taken on a Zeiss Cell Observer SD (Carl Zeiss, Oberkochen, Germany) using a 20x objective. Analysis of live and dead cell count was manually performed with ImageJ (NIH).

### 2.7 Thiol quantification

The Measure-iT thiol kit was used to quantify unreacted thiols (ThermoFisher). Di-functional peptides or PEG dithiol were reacted with PEG-maleimide in 10 µL volumes for 10 minutes before reacting with 100 µL of the Measure-iT thiol working solution.

### 2.8 Bead dispersion and image analysis

Submicron-sized fluorescent particles (Blue 0.5µm Fluoro-Max, Thermo) were embedded within the hydrogel. The appropriate amount of beads for a 1g/L suspension in the hydrogel are pelleted, the supernatant removed, and then resuspended in the bulk polymer macromere suspension. Hydrogels are formed on a glass bottom well-plate (no. 1.5 coverslip glass; In Vitro Scientific, Sunnyvale, CA) that was plasma treated and subsequently thiol-silanized with 2 vol% (3-mercaptopropyl)trimethoxysilane (Thermo) in 95% ethanol (pH 5). Hydrogels were suspended in 1x PBS at pH 7.4 and allowed to swell overnight. Fluorescent images were taken on a Zeiss Cell Observer SD (Carl Zeiss) suing a 40x oil immersion objective. Analysis of bead density was manually performed in ImageJ.

### 2.9 Small molecule diffusion

Bulk diffusion of Rhodamine 6G (R6G) (Stokes’ radius 0.76 nm, Sigma-Aldrich) in hydrogels was measured by encapsulating 0.1 g/L R6G in the hydrogel and sampling the supernatant at 5-minute intervals for 2 hours. The samples were analyzed on a fluorescent plate reader at an excitation/emission of 526/555nm. The diffusion coefficient of R6G was calculated using the modified Fick’s law for solute release behavior of swelling systems

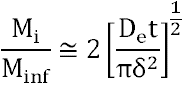

where *M*_*i*_ is the concentration of released solute at time *i*, *M*_*inf*_ is the solute concentration at infinite time, *D*_*e*_ is the effective diffusion, *t* is time, and δ is half of the hydrogel thickness. The mass balance to calculate *M*_*i*_ is

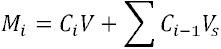

where *C*_*i*_ is the released solute concentration of the solute at time *i*, *V* is the volume of the bulk solution, and *V*_*s*_ is the volume of the sample[25].

### 2.10 Statistical analysis

Data are reported as the mean ± standard deviation. Significance analyses were performed using GraphPad Prism 7.0 software. Unless otherwise noted, statistical significance was determined using a two-tailed t-test and p-values <0.05 were considered significant, where p<0.05 is denoted with *, ≤0.01 with **, ≤0.001 with ***, and ≤0.0001 with ****.

## 3. Results and Discussion

### 3.1 Weak bases sufficiently catalyze the thiol-maleimide reaction

With the goal of creating PEG-based hydrogels via the thiol-maleimide Michael addition with homogeneous networks, we explored several mechanisms to slow down this typically very fast reaction. Triethanolamine (TEOA) is a strong base commonly used to increase thiolate formation under physiological pH [8,17]. We hypothesized a weaker base would decrease the gelation speed by reducing the accumulation of thiolate groups. We coupled thiolate groups across the double bond of the maleimide in the presence of bases of differing catalytic strength, while simultaneously varying the overall polymer wt% (**Figure 1a**). The hydrogen phosphate in phosphate buffered saline (PBS, pH 7.4) sufficiently catalyzed the hydrogel formation reaction without TEOA (**Figure 1b**). The material bulk modulus and the percentage of unreacted thiols also did not change significantly; suggesting the mechanism of gelation is conserved (**Figure 1b-c, Supplemental Figure 1**). Interestingly, the reaction speed was not dramatically different between PBS and TEOA. TEOA increases the speed of thiolate formation, which could correlate with polymerization speed. Since an increase in polymerization speed was not observed, we suggest that in PBS the thiolate formation is already in excess.

**Figure 1.**
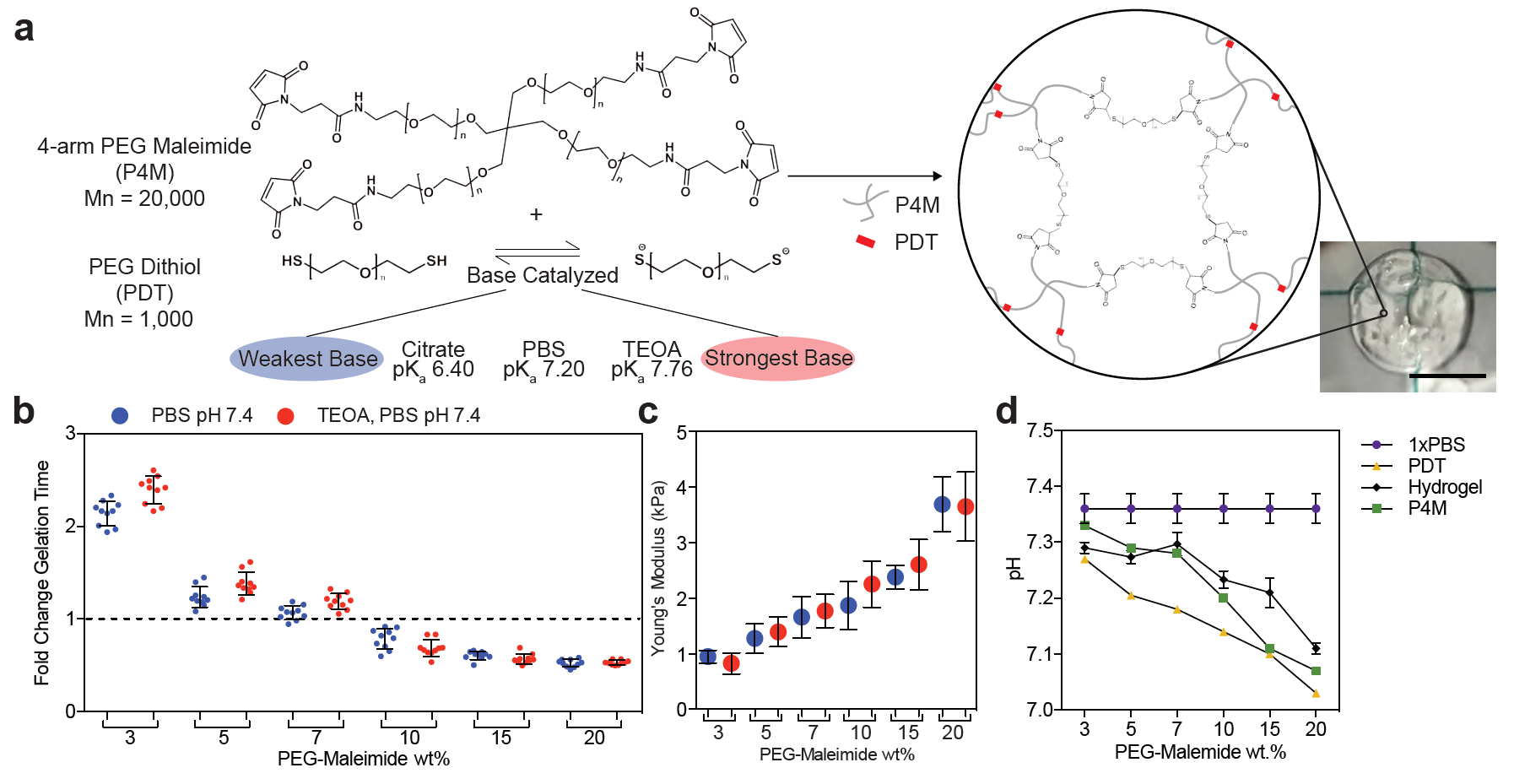
A weak base catalyzes the thiol-maleimide reaction. a) Schematic of the Michael-type addition reaction for a 4-arm PEG-maleimide (P4M) and linear PEG-dithiol (PDT) hydrogel. Thiolates catalyzed by a base initiate the reaction with a nucleophilic addition onto the alkene group in the maleimide, forming a stable bond and assembling the polymeric network. Image of a resulting 10µL 20wt% hydrogel made in PBS at pH 7.4 (right, scale 5mm). b) Fold change in hydrogel gelation time in two buffers: PBS (blue) and 2mM TEOA in PBS (red). The dashed line marks the internal control. c) Young’s modulus and d) pH for the hydrogel and precursors versus polymer wt%. The lines connect data from the same condition and do not represent a model fit. Error bars represent the SD, N≥3.

Decreasing the overall polymer wt% slowed gelation time (**Figure 1b**). This result happened even though the pH of the precursor solutions decreased with increasing polymer wt%, which should lower the ability of the thiol to react (**Figure 1d**). This indicates that lower concentrations of reactive groups slow gelation more effectively than the small pH change observed [18]. However, even at low polymer wt% and in a weaker base, hydrogels polymerized within 5 seconds, which is comparable to the speeds others have reported [14]. This is still faster than most users, or automated systems, could uniformly mix the precursor solutions, and the resulting hydrogels are visibly wrinkled (**Figure 1a**). Tuning substrate stiffness is also important for many biological studies, making it necessary to find approaches that reduce the hydrogel reaction kinetics across a range of polymer and crosslinker wt%.

### 3.2 The pKa of the Michael-type donor regulates the speed of the thiol-maleimide reaction

Our ability to form these hydrogels without the TEOA catalyst led us to explore whether even weaker bases and buffer capacity could support the thiol-maleimide reaction. Buffering capacity depends on the strength of the conjugate base formed at a given pH. While others have mainly modulated the pH to slow this reaction, we compared the effect of both the pH and the strength of the conjugate base. At pH values between 5.8-7.4, the conjugate base is hydrogen phosphate in PBS, and sodium citrate in citrate (**Figure 2a**). The thiol-maleimide reaction is reliant on the thiol pKa, because the thiol/thiolate equilibrium is modulated by buffer pH, as described by the Henderson-Hasselbalch relationship [14,17,18]. We explored pHs between 5.8 and 10 because this is the range where physiological reactions occur [26]. Reducing the strength of the conjugate base when switching from PBS to citrate decreased gelation speed, but changing the solution pH had a stronger effect (**Figure 2b**). As expected, more basic pHs increased the reaction speed, while acidic pHs slowed the reaction.

**Figure 2.**
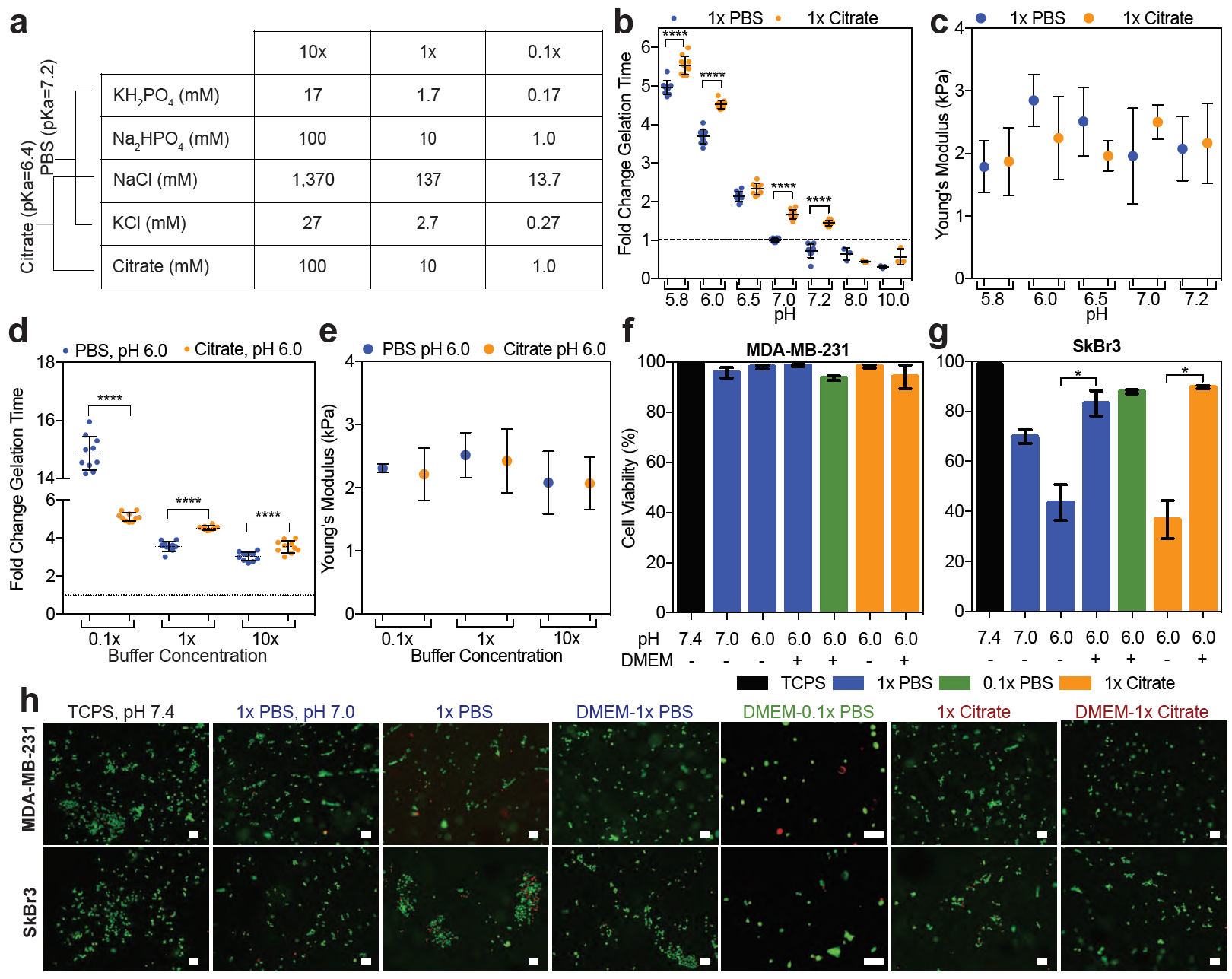
Buffer concentration and pH control the thiol-maleimide reaction rate without altering hydrogel modulus. a) Salt concentrations in PBS and citrate buffers. b) Fold change in gelation time for hydrogels formed in 1x PBS (blue) and 1x citrate (yellow) at different initial pHs. The dashed line marks the internal control. c) Young’s modulus of hydrogels with respect to buffer and pH. d) Fold change in gelation time and e) the Young’s modulus of resulting hydrogels versus buffer concentration. Error bars represent the SD, N≥3. A live (green) and dead (red) stain used to assess percent cell viability for f) MDA-MB-231 and g) SkBr3 cells encapsulated in hydrogels with PBS at pH 7.0 or PBS and citrate buffer at pH 6.0 in the presence or absence of the high glucose supplement, DMEM, 2 hours post-encapsulation. . h) Representative images of the stain. Unless stated all buffers are 1x. Error bars represent the SD, N≥3.

The pH was stable between the precursor polymer solutions and post-gelation, which is important for cell encapsulation (**Supplemental Figure 2a-b**). Additionally, changing the buffer did not significantly change the hydrogel bulk modulus (**Figure 2c**). At the lowest pH tested, a dilute PBS buffer (0.1x) dramatically increased the gelation time without changing hydrogel pH or modulus (**Figure 2d-e, Supplemental Figure 1c**). This result could be because at pH 6.0, hydrogen phosphate equilibrium favors the formation of its conjugate acid in the 0.1x PBS to maintain a constant pH (**Supplemental Figure 2c**), thereby reducing thiolate formation. This was only observed with PBS, because a pH of 6.0 is closer to the pKa of citrate (6.4) than of PBS (7.2).

### 3.3 High glucose medium maintains cell viability in acidic gel polymerization buffers

Although a pH of 6 is within the physiological range for many biochemical reactions, we were worried this condition would decrease cell viability. In previous work, people have used fibroblasts to screen the impact of changing hydrogel conditions on cell viability[14]. However, fibroblasts are largely insensitive to drastic changes in pH and serum, likely because their role in wound healing exposes them to a variety of environmental conditions [27]. Thus, we used the MDA-MB-231 and the SkBr3 breast cancer cell lines because they have been used in high-throughput drug screening [28,29], a potential application for these materials, and it is unknown how sensitive they are to external buffer changes. We focused on the conditions that most effectively reduced the reaction speed: PBS and citrate at low pH. Buffers were tested at pH 6.0 in the presence or absence of the high glucose supplement, DMEM, and cell viability was compared to encapsulation in PBS at pH 7 or culturing cells on tissue culture polystyrene (TCPS) (**Figure 2f-g**).

Across all tested buffer conditions, the MDA-MB-231 cells were nearly 100% viable 2 hours post-encapsulation (**Figure 2f**). The SkBr3 cells were very sensitive to the encapsulation conditions, but their viability increased when the buffer solution was mixed with a high glucose supplement (**Figure 4g**). The addition of the high glucose supplement did not influence the gelation time of these hydrogels (**Supplemental Figure 2d**). Since acidic environments have been shown to stimulate tumor invasion [30], this may explain the pH insensitivity observed in the metastatic MDA-MB-231 cells and the pH sensitivity in the minimally tumorigenic SkBr3 cells.

### 3.4 Electronegative crosslinkers effectively slow gelation speed while maintaining high cell viability

Others have used negatively charged amino acids near the thiol to slow the reaction kinetics by increasing the thiol pKa through electrostatic interactions [14,19]. Most of these electronegative peptides sequences are included in hydrogels to facilitate cell degradability [20]. Though a large number of peptides have been shown to be susceptible to degradation by cells, only one of these sequences has been used to study the kinetics of polymer network formation [14]. We explored a panel of peptides with biological function to see how the different amino acid sequence combinations change the polymerization rate. Though all of these peptides had a negatively charged aspartate near the thiol, we only observed a drastic change to gelation speed with the IPESLRAG sequence (**Figure 3a**). This peptide contains a glutamate in the degradable sequence, and glutamates have been shown to reduce the speed of this reaction better than aspartates [14].

**Figure 3.**
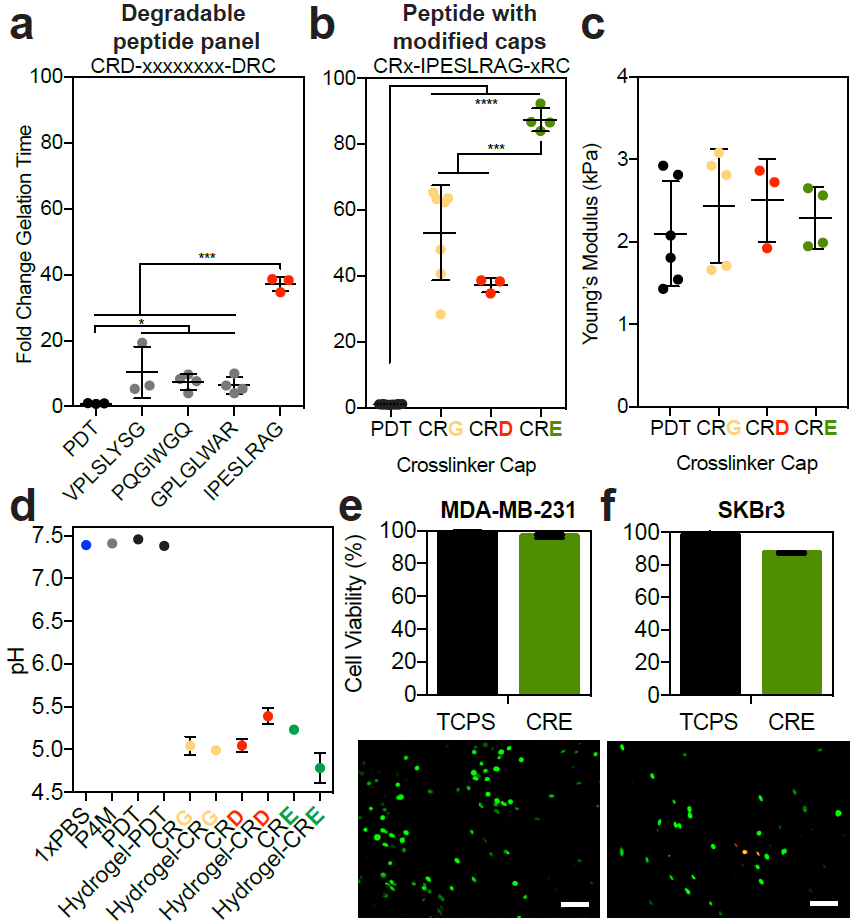
Increasing crosslinker electronegativity slows the reaction. Different dithiol crosslinkers and modifications around the thiol were used to form a hydrogel: PDT (black) and crosslinkers with an aspartate (CRD, gray or red), a glycine (CRG, yellow), or a glutamate (CRE, green) near the cysteine. a) The fold change in gelation time for a series of degradable peptides with an aspartate near the cysteine, where the sequence is the x-axis label. b) The fold change in gelation time for the top performing peptide with different amino acids directly adjacent to the reactive cysteine (“modified caps”). c) Young’s modulus for resulting hydrogels, and d) pH of each precursor solution (P4M: red, PBS: blue) for some of the crosslinkers. Error bars represent the SD, N≥3. Error bars are not visible in conditions with low variation. A live (green) and dead (red) stain used to assess percent cell viability for e) MDA-MB-231 and e) SkBr3 cells encapsulated in hydrogels with the CRE crosslinker at in 1X PBS at pH 7.4, 2 hours post-encapsulation. Below each graph are representative images of the stain. Error bars represent the SD, N≥3.

We then decided to modify the cap (the amino acid adjacent to the reactive cysteine) to include either a glutamate or glycine, instead of an aspartate. None of these amino acid substitutions altered the final bulk hydrogel modulus, and including a glutamate adjacent to the thiol was the most effective way to slow the reaction (**Figure 3a-c**), consistent with work done by others [14]. These amino acid substitutions in the crosslinker lowered the precursor solution to a pH of ∼5.8 (**Figure 3d**), so we thought it possible that the pH changes were responsible for the gelation time differences observed. However, manual adjustments to this pH only increased the gelation time by ∼6-fold (**Figure 2b**), and therefore could not have accounted for this extent of slowing (Figure 3d shows 40-100 fold changes). As shown here, and by others [14,19], modulating the pKa of the thiol via electronegative crosslinkers is the most effective way to slow gelation. However, this is the first thorough characterization of a panel of degradable peptides to modulate the thiol-maleimide reaction speed. Lastly, we wanted to ensure that cells were viable under these reaction conditionsand found that viability for both the MDA-MB-231 and SkBr3 cell lines in the hydrogel crosslinked with the electronegative peptide remained above 80% (**Figure 3e-f**). We speculate this high viability is likely because the encapsulation conditions are at an optimal cell culturing pH of 7.4.

### 3.5 Hydrogel uniformity impacts diffusion and particle distribution

We next sought to explore how the speed of gelation could impact the implementation of thiol-maleimide hydrogels into applications that would require large amounts of consistently formed hydrogels, such as high throughput screening. We chose four conditions that had varying speeds of gelation: 1x PBS at pH 7.0, 0.1x PBS at pH 6.0, and crosslinked with a CRE or CRG cap near the thiol at pH 7.4 (polymerization conditions listed from fasted to slowest gelation times). Though the bulk stiffness of these materials is not different (**Figure 2c,e, Figure 3c**), it is clear from visual inspection that slower reaction speeds create more uniform materials (**Figure 4a**). The number of unreacted thiols remained constant across these reaction conditions (**Figure 4b**), thus the number of possible crosslinks does not influence this heterogeneity. Others have also reported these observable gel wrinkles, which they attributed to local differences in crosslinking [14], and our data shows that these densities do not influence the total number of thiols that react in these materials.

**Figure 4.**
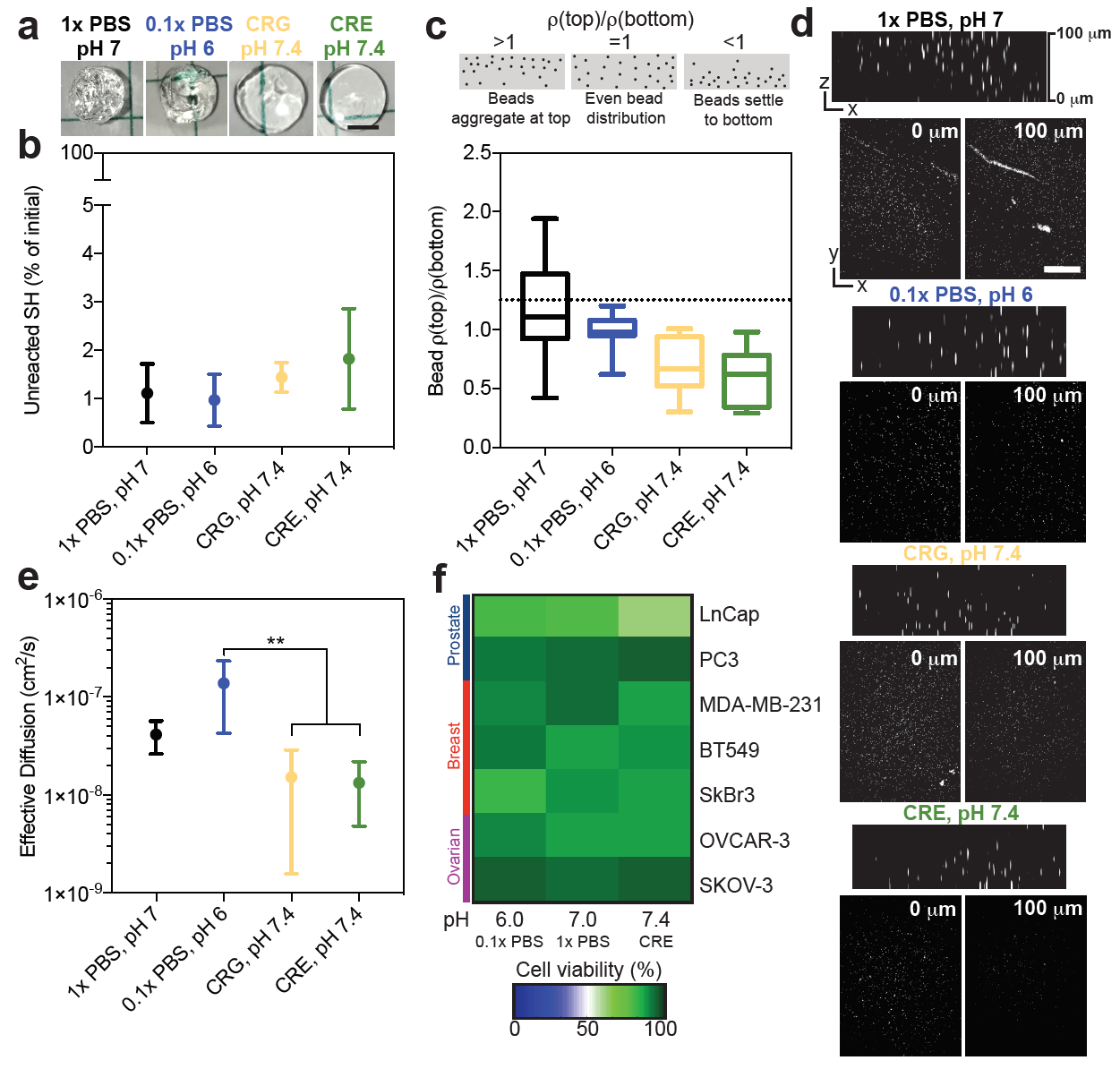
Optimizing kinetics for hydrogel applications. Four reaction conditions were studied further: 1x PBS, pH7 (black), 0.1x PBS, pH 6 (blue), hydrogel crosslinked with a CRG (yellow) or CRE (green) peptide in PBS at pH 7.4. They are ordered for slow (left) to fast (right) gelation speed. a) Representative images for select hydrogels, pre-swelling, formed at the different reaction conditions. b) Percentage of unreacted thiols from initial versus reaction condition. c) Fluorescent bead density in hydrogels at 100µm (top) divided by density at 0µm (bottom) compared to reaction condition. d) Representative images of the beads at 0µm and 100µm. Error bars represent the SD, N=4, n=4. e) Effective diffusion of R6G through hydrogels formed at different reaction conditions. f) A live and dead stain was used to assess percent cell viability for different cancer cell lines encapsulated in hydrogels with differing reaction conditions conditions in the presence or absence of the high glucose supplement, DMEM, 2 hours post-encapsulation.

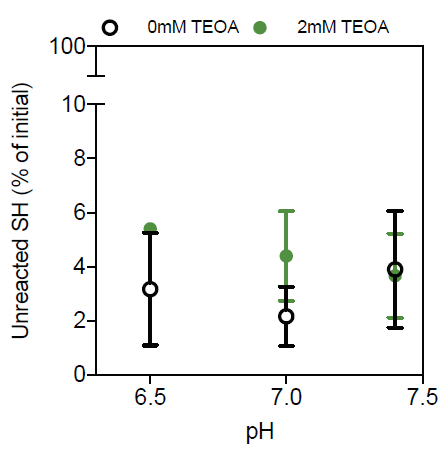

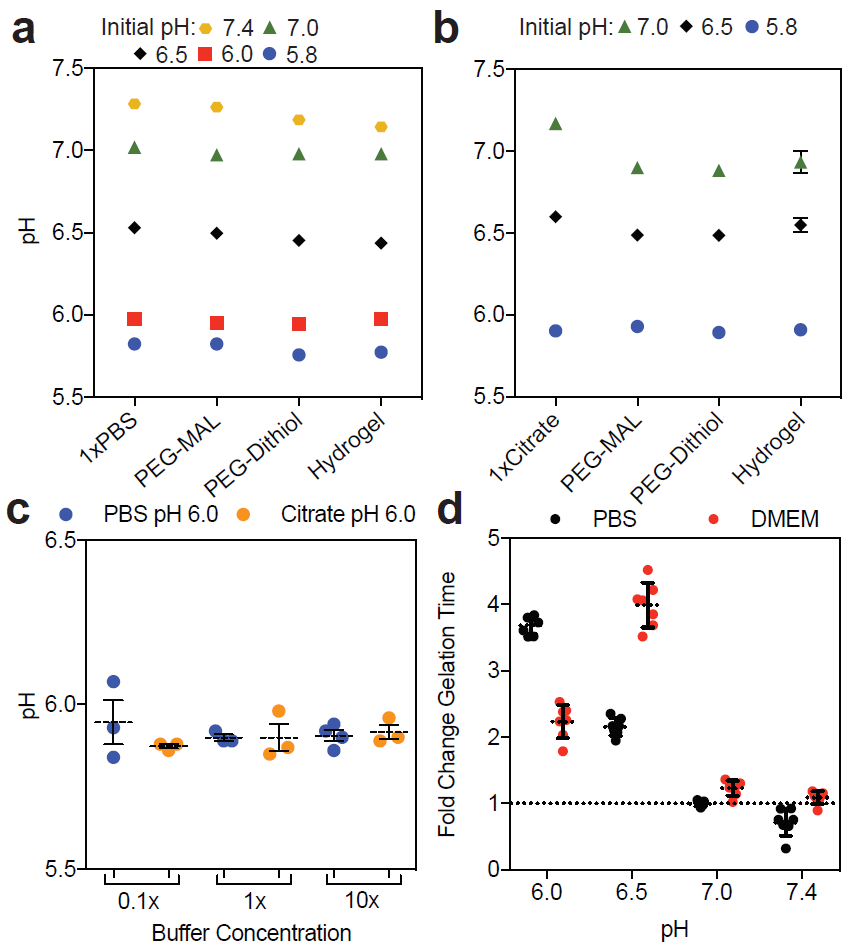

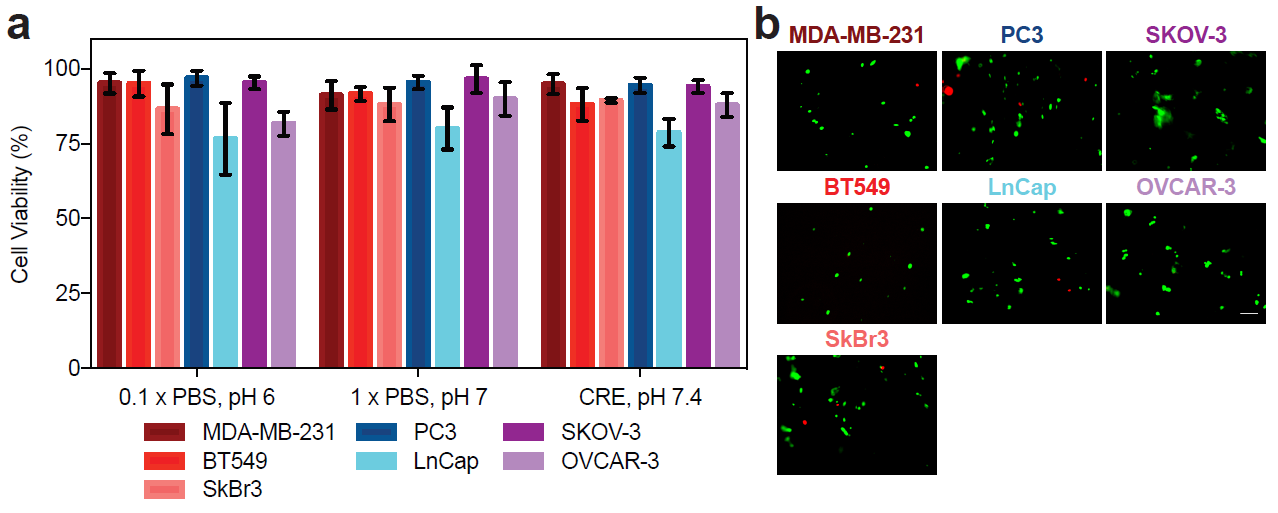

One potential consequence of polymer density gradients is a non-uniform distribution of cells during encapsulation. We explored this by encapsulating fluorescent beads into hydrogels formed under varying conditions. We quantified the bead density in images acquired throughout the bulk of the gel (bottom, near objective=0µm, and top=100 µm). Forming hydrogels in 0.1x PBS at pH 6.0 resulted in the most uniform particle distribution (**Figure 4c-d**). The fastest polymerization conditions (1x PBS at pH 7.0) had the largest amount of variability in bead distribution, indicating that the reaction formed too quickly and resulted in massive gel polymerization heterogeneities (**Figure 4d**). Conversely, hydrogels formed slowly with the electronegative peptide crosslinkers had a higher bead density near the bottom of the hydrogel than the top, likely because gelation was too slow and the beads settled before the network was fully formed. Overall, this indicates that the optimal gelation needed to achieve uniform particle distribution and minimally heterogeneous gels is approximately 30 seconds, and is achieved with the low ionic strength buffer at slightly acidic pH.

Though the polymer densities did not change the number of thiols reacted or the bulk modulus of the final hydrogel, we speculated that it would influence the diffusion of small molecules. Diffusion of R6G, a molecule with a molecular weight comparable to many drugs of interest for cancer applications and regenerative medicine, was significantly faster through hydrogels formed with visible heterogeneities (**Figure 4e**). We attribute this to the areas of un-polymerized network, resulting in local void spaces where the drug can immediately diffuse out. This could limit the use of these materials in drug screening applications, because nonuniform gradients of drugs will be presented to cells in some areas. However, while the slower conditions had less variable diffusion, the distribution of large particles was heterogeneous (**Figure 4c**), so both these factors must be taken into account when optimizing gelation speed for a specific application of interest.

Finally, we quantified how these differences in the hydrogel reaction conditions impacted cell viability for cell lines that we have previously used for drug screening applications with this hydrogel platform [28]. We measured the viability of three breast cancer (MDA-MB-231, SkBr3, and BT549), two prostate (PC3 and LnCap), and one ovarian cancer cell line (OVCAR-3 and SKOV-3) across these same gelation conditions (**Figure 4f, Supplemental Figure 3**). Certain cell lines were particularly sensitive to the reaction conditions (LnCap, SkBr3, and OVCAR-3), but we did not observe a consistent sensitivity across any cancer type. Somewhat surprising to us, the most sensitive cell lines (LnCap and SkBr3) were least viable in opposing conditions. The LnCap cells were not very viable in when encapsulated with the electronegative crosslinkers, and the SkBr3s were very sensitive to pH. Although it is possible that these reaction conditions would not change the long-term response of cells to the hydrogel, such as proliferation, motility, etc., this is a critical consideration for groups that want to use a single gel platform to compare across many cell types. Cell proliferation is sensitive to starting cell numbers, and this is not consistent across the gelation conditions and cell lines studied here. We recommend that groups using this hydrogel system consider the optimal gelation time, coupled with cell sensitivity to hydrogel polymerization conditions for their application of interest.

## 4. Conclusion

The PEG-maleimide hydrogel is very attractive in tissue engineering and regenerative medicine because of its high crosslinking efficiency, biocompatibility, and ability to functionalize with bio-active peptide groups. A major drawback of this system is that the fast gelation speed can result in documented crosslinking heterogeneities. We compared how catalytic buffer strength, pH, and electronegative crosslinkers controlled the thiol-maleimide reaction while retaining high cell viability without changing final bulk modulus. Certain cell lines were sensitive to the reaction conditions, and high glucose preserved cell viability, an important consideration for users encapsulating cells. We also confirmed the results of others that coupling a glutamate near the cysteine of the peptide crosslinker slowed the thiol-maleimide reaction by 90-fold. We add that reducing the pH and ionic strength of the buffer was the most efficient way to slow the reaction without chemical modifications to the crosslinker. These adaptations influenced the uniformity of particle dispersion and small molecule diffusion through the matrix. Overall, although the slowest reaction speeds led to the most uniform hydrogels, the reaction speeds were too slow to ensure that particles did not settle during gelation. Therefore, we suggest that users consider these factors when creating thiol-maleimide hydrogels more consistently and simultaneously amenable to applications of interest.

## Acknowledgements

We would like to acknowledge Thomas McCarthy for his technical assistance in peptide synthesis. We would like to thank Dr. Sarah Perry, Dr. Lila Gierasch, and Dr. Alfred Crosby for use of equipment. Research reported in this publication was supported by the National Institute of Health Office of the Director under Award Number S10OD010645. SRP is a Pew Biomedical Scholar supported by the Pew Charitable Trusts. SRP was supported by a faculty development award from Barry and Afsaneh Siadat. This work was funded by an NIH New Innovator award (1DP2CA186573-01) and an NSF CAREER award (DMR-1454806) to SRP.

